# Platypus: an open-access software for integrating lymphocyte single-cell immune repertoires with transcriptomes

**DOI:** 10.1101/2020.11.09.374280

**Authors:** Alexander Yermanos, Andreas Agrafiotis, Josephine Yates, Chrysa Papadopoulou, Damiano Robbiani, Florian Bieberich, Rodrigo Vazquez-Lombardi, Daniel Neumeuer, Annette Oxenius, Sai T. Reddy

## Abstract

High-throughput single-cell sequencing (scSeq) technologies are revolutionizing the ability to molecularly profile B and T lymphocytes by offering the opportunity to simultaneously obtain information on adaptive immune receptor repertoires (VDJ repertoires) and transcriptomes. An integrated quantification of immune repertoire parameters such as germline gene usage, clonal expansion, somatic hypermutation and transcriptional states opens up new possibilities for the high-resolution analysis of lymphocytes and the inference of antigen-specificity. While multiple tools now exist to investigate gene expression profiles from scSeq of transcriptomes, there is a lack of software dedicated to single-cell immune repertoires. Here, we present Platypus, an open-source software platform providing a user-friendly interface to investigate B cell receptor (BCR) and T cell receptor (TCR) repertoires from single-cell sequencing experiments. Platypus provides a framework to automate and ease the analysis of single-cell immune repertoires while also incorporating transcriptional information involving unsupervised clustering, gene expression, and gene ontology. To showcase the capabilities of Platypus, we use it to analyze and visualize single-cell immune repertoires and transcriptomes from B and T cells from convalescent COVID-19 patients, revealing unique insight into the repertoire features and transcriptional profiles of clonally expanded lymphocytes. Platypus will expedite progress by increasing accessibility to the broader immunology community by facilitating the analysis of single-cell immune repertoire and transcriptome sequencing.

## Introduction

Immune repertoires are comprised of a diverse collection of BCRs and TCRs, which enable molecular recognition to a vast number of pathogen and disease antigens. Immune repertoire diversity is initially generated as a result of lymphocyte V(D)J recombination and, in the case of B cells, can undergo further sequence diversification in the form of somatic hypermutation. Targeted deep sequencing of BCRs and TCRs from bulk populations of lymphocytes has paved the way to quantify the diversity, distribution and evolution of immune repertoires ^1–4^. However, one major challenge in immune repertoire sequencing is acquiring information on correct receptor chain pairing [variable light (VL) and variable heavy (VH) for BCR and variable alpha (Vα) and variable beta (Vβ) for TCR], which greatly complicates identification of clonal groups and antigen-specificity ^5,6^. Until only recently it was not possible to directly integrate a lymphocyte’s phenotypic gene expression information (i.e., activation, exhaustion, antibody secretion) with its immune receptor sequence.

Recent developments in microfluidic and scSeq technologies have now made it possible to obtain information on immune repertoires or transcriptional profiles at high-throughput ^7–9^. Several of these methods have been tailored specifically for lymphocytes, thus making it possible to perform parallel sequencing of immune repertoires and whole transcriptomes ^10,11^. Furthermore, commercially available instruments and protocols [10X Genomics Chromium and VDJ and gene expression (GEX) libraries] are further accelerating progress in this field. This simultaneous sequencing of immune repertoires and transcriptomes produces single-cell datasets with features ranging from quantitative gene profiles, cellular phenotypes, transcriptional clustering, clonal diversity and expansion, germline gene usage, somatic hypermutation, among many others. These high-dimensional datasets can be mined to discover novel insight on lymphocyte immunobiology, function, and specificity. For example, one recent study leveraged scSeq to discover the distinct transcriptional profiles and specificity of B cells following influenza vaccination ^8^. In another study, clonal expansion and activation signatures of tumor-infiltrating T cells were profiled ^12^. Other studies have leveraged this technology to answer fundamental questions across a variety of areas in immunology such as bacterial infection responses ^13^, tumor-immune microenvironment ^14^, clonal expansion in Alzheimer’s disease ^9^, and B cell differentiation ^15^.

While multiple bioinformatic tools exist to facilitate rapid analysis of gene expression from scSeq ^16–19^, they do not allow the incorporation of immune repertoire information. Analogously, existing software packages to analyze immune repertoires do not allow the user to supply accompanying gene expression and transcriptome data ^20–22^. Taken together, these considerations complicate the analysis of BCR and TCR repertoires for those with little bioinformatics experience and who are unfamiliar with the output data from scSeq experiments. To address the lack of software specifically tailored to single-cell lymphocyte sequencing data, we developed Platypus, an open-source R package that contains an automated pipeline to analyze and integrate single-cell immune repertoires with transcriptome data. With only a few lines of code, Platypus allows users to easily extract immune repertoire features such as clonal expansion, somatic hypermutation, isotype switching and integrate it with transcriptome features such as differential gene expression. We subsequently demonstrate the value of this package using scSeq data from convalescent coronavirus infectious disease 2019 (COVID-19) patients. Our analysis revealed clonal expansion in B and T cells, and within these we could identify distinct patterns of somatic hypermutation, amino acid usage, clonal convergence and transcriptional heterogeneity. Taken together, Platypus helps facilitate the analysis of single cell immune repertoires and transcriptomes and reveal novel insights such as the transcriptional profile of clonal expanded and potentially pathogen-reactive lymphocytes.

## Results

### Single-cell immune repertoire sequencing analysis

Platypus allows the user to integrate single-cell immune repertoire and transcriptome sequencing data, which includes automation of pre-processing, filtering, and data visualization (Figure 1). While Platypus is optimized for data generated by the 10X Genomics System, it is also adaptable to other cell-barcode based scSeq data (e.g. RAGE-seq, Split-Seq ^10,23^). Users can simply supply the path to the output directory from the 10X cellranger analysis as input to Platypus, which then extracts and annotates key immune repertoire metrics such as clonal diversity, clonal expansion, somatic hypermutation, reference germline gene usage, and sequence motifs (Figure 1). Platypus can perform additional clonotyping, either increasing or relaxing the pre-determined stringency of upstream alignment tools by incorporating information regarding germline gene usage or sequence homology thresholds. In addition to clonal sequence information [based on complementarity determining region 3 (CDR3)], it also extracts full-length sequences of both immune receptor variable chains (VH and VL for BCRs and Vα and Vβ for TCRs). Furthermore, Platypus enables the quantification, comparison, and visualization of more advanced repertoire features such as sequence similarity networks ^1^, phylogenetic tree construction ^24,25^, isotype quantification, and diversity metrics ^26^.

**Figure 1.**
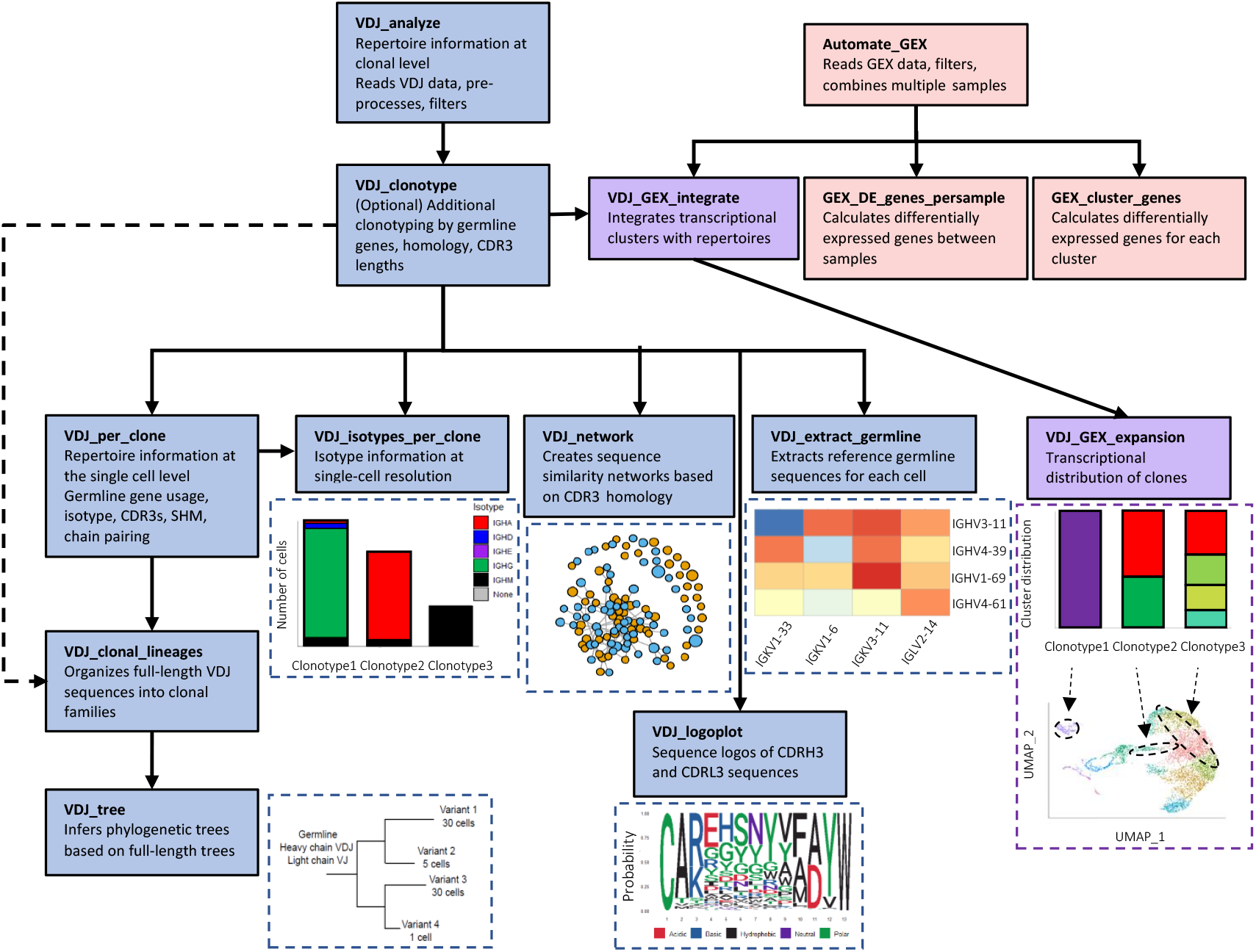
Flowchart demonstrating the workflow of Platypus. A selection of functions internal to Platypus and their respective relationship are depicted. Each node in the flow chart indicates a process in the workflow requiring just a single line of code with user-definable parameters.

To demonstrate the potential of Platypus, we performed single-cell immune repertoire and transcriptome sequencing on B and T cells isolated from peripheral blood mononuclear cells (PBMCs) of two convalescent COVID-19 patients (Figure 2A). 10X Genomics’ basic alignment tool is cellranger, which has a default clonotyping strategy that groups identical CDRH3 + CDRL3 nucleotide sequences into clonal families; this approach would be too restrictive in identifying clonotypes of B cells that have undergone somatic hypermutation in the CDR3s. We thereby demonstrated the ability and impact of changing the clonotyping strategy to include germline genes, CDR3 length restrictions, and sequence homology requirements for the B cell repertoires of the two COVID-19 patients, which resulted in a decrease in the number of unique clones when additional repertoire features were included in the clonotyping definition (Figure 2B). Next, using Platypus, we were able to detect and visualize clonal expansion for both B and T cells (Figures 2C, 2D). Furthermore, we were able to relate isotype information with clonal expansion at single-cell resolution, thereby observing that the majority of most expanded B cell clones were of the IgA isotype (Figure 2D). We next used built-in functions of Platypus to extract other common immune repertoire statistics and features, such as CDR3 length distribution and common sequence space motifs (sequence logo plots). This revealed tremendous diversity in the B cell response at the most common paired CDRH3 + CDRL3 amino acid sequence length in both COVID-19 patients (Figures 2E, 2F). While we focused on the most frequent CDRH3 + CDRL3 sequence length, such an analysis using Platypus could theoretically be applied to other single-cell subsets, such as those cells with known antigen-specificity or those B cells that underwent extensive somatic hypermutation.

**Figure 2.**
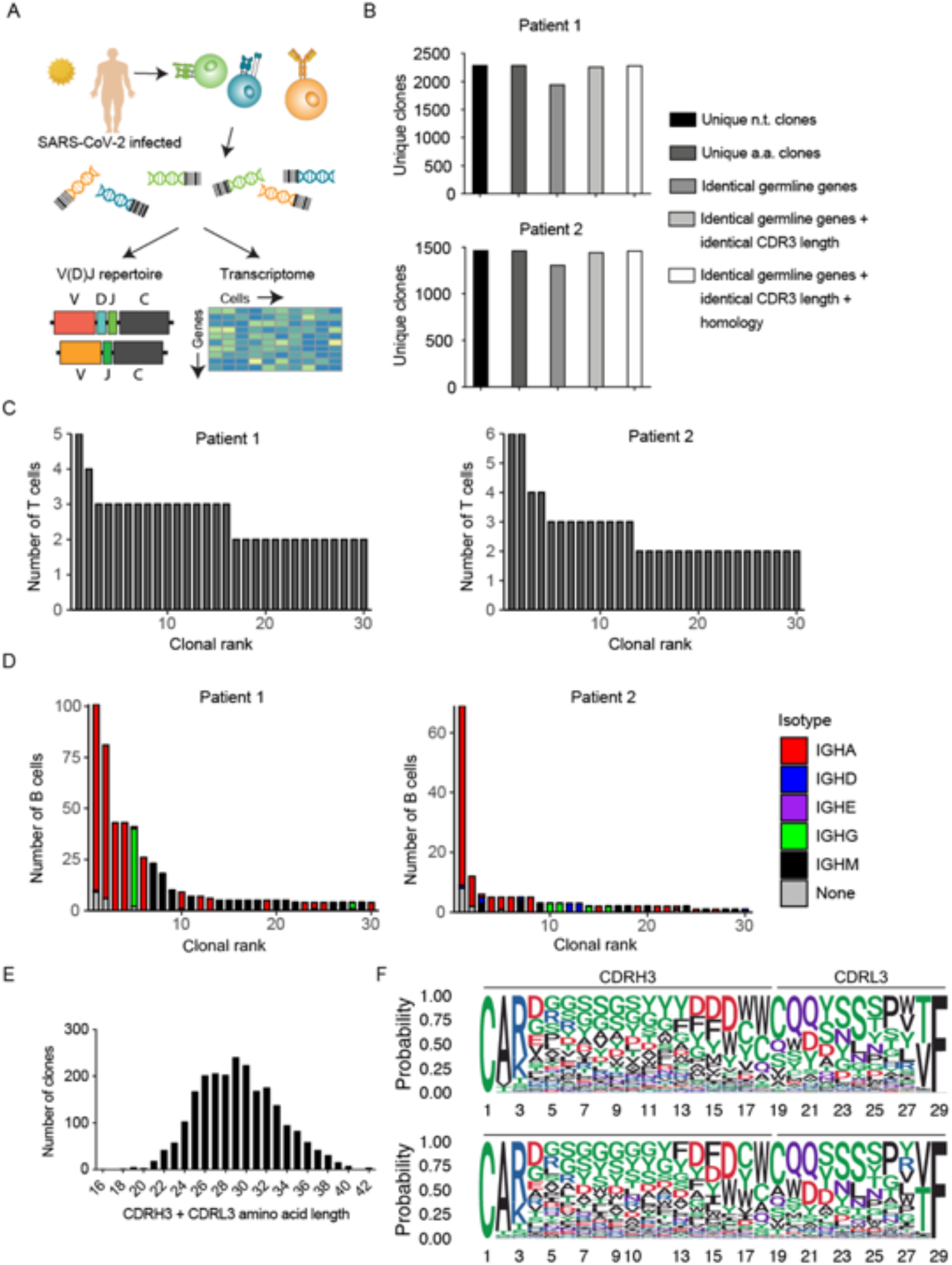
Extracting and visualizing clonal information of PBMCs from patients recently infected with SARS-CoV-2 using Platypus. (A) Experimental overview for single-cell immune repertoire of B and T cells in two patients previously infected with SARS-CoV-2. (B) Multiple B cell clonotyping strategies involving CDR3 sequence identity, germline gene usage, and sequence homology thresholds from the two patients previously infected with SARS-CoV-2. (C) Clonal expansion profiles of the T cells from the blood repertoires of two individuals recently infected with SARS-CoV-2. Clone is defined as unique CDRβ3 + CDRα3 nucleotide sequence. (D) Clonal expansion profiles of the T cells from the blood repertoires of two individuals recently infected with SARS-CoV-2. Clone is defined by unique CDRH3 + CDRL3 nucleotide sequence. Color depicts isotype on the cell level within each clone determined by the VDJ repertoire sequencing libraries. All functions and plots demonstrated require less than five lines of code. (E) Length distribution of the paired CDRH3 + CDRL3 amino acid sequences from the B cell clones of a single patient. (F) Sequence logo plots of those paired CDRH3 + CDRL3 amino acid sequences with a combined sequence length of 29 in two patients. Colors correspond to biochemical properties: red=acidic, blue=basic, black=hydrophobic, purple=neutral, green=polar. Top logo plot corresponds to patient 1 and bottom logo plot corresponds to patient 2.

### Integration of single-cell immune repertoires and transcriptomes

A critical feature of Platypus is that it can seamlessly integrate single-cell immune repertoire data with transcriptome sequencing data. It allows users to directly interact with the commonly used scSeq transcriptome analysis program Seurat ^16^, while tuning parameters specifically relevant for immune repertoires. Therefore, we next investigated additional repertoire and transcriptional data of highly expanded B and T cell clonal groups, which allows us to relate repertoire information (e.g. expansion, CDR3 sequence, isotype) to phenotypic cellular behavior (e.g. whether a cell is proliferating, differentiated, activated, exhausted, etc.). We first integrated the transcriptome sequencing data from both COVID-19 patients by normalizing and scaling the data using default parameters in Seurat (although Platypus supports other normalization methods such as SCTransform and Harmony) ^16,17^. We could thereby compute clusters based on gene expression and subsequently visualize the cells from each patient on the same two-dimensional uniform manifold approximation projection (UMAP) plot (Figure 3A). Overlaying the gene expression of CD4, CD8, CD3, and CD19 revealed a separation between B and T cells, in addition to demonstrating similarly distributed lymphocytes populations arising from both patients (Figure S1A). Of note is that we detected CD3 expression in the B cell clusters on the UMAP, in addition to minor CD19 expression in the T cell clusters (Figures 3A, S1A), together suggesting the possibility off doublets in which B and T cells were present in the same microfluidic droplet. We next used Platypus to investigate which transcriptional-based cluster contained the most expanded T cell clonal families. We could demonstrate in one of the COVID-19 patients that some of the most expanded T cell clones were located across multiple transcriptional clusters, with 6 of the most expanded clones located in the CD8+ T cell clusters and the remaining 4 were located in the CD4+ T cell clusters (Figures 3B, S1A). Furthermore, we observed that the majority of expanded CD8 T cells were located in cluster 6, which corresponded to high expression of granzymes and perforin (Figure S1B). Another option to visualize transcriptional heterogeneity using Platypus is to overlay clonal information directly onto the UMAP plot. To investigate the transcriptional heterogeneity of clonally expanded T cells, we supplied the clonal index of the top 10 most expanded clones from a single patient and visualized where these cells lie within the UMAP plot, again revealing that many of the expanded clones were located in cluster 6 (Figure 3C). This approach can be leveraged to profile transcriptome signatures of well-defined clones, for example ones in which antigen-specificity is known.

**Figure 3.**
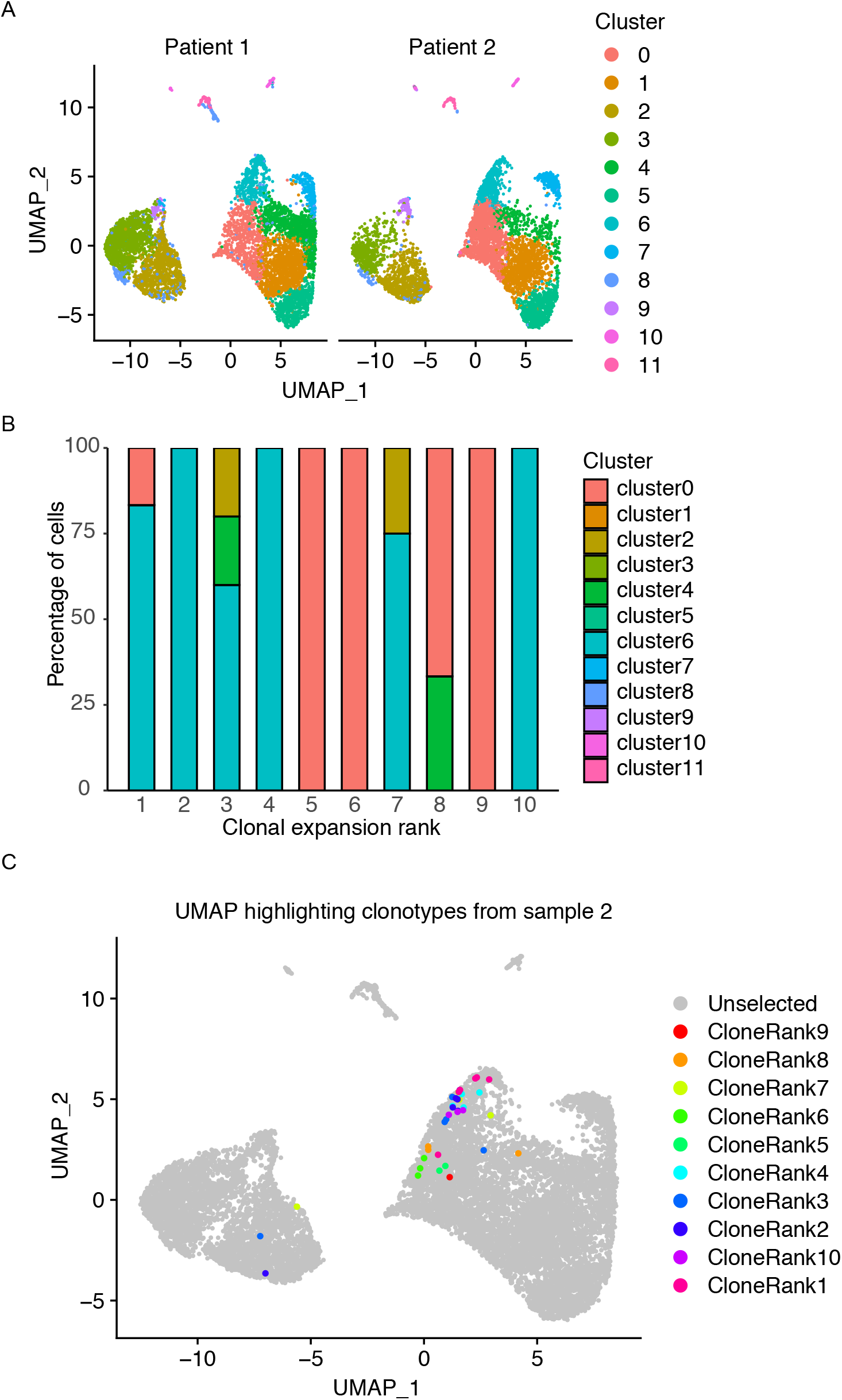
Integration of gene expression (GEX) and repertoire (VDJ) sequencing datasets from two patients recently infected with SARS-CoV-2. (A) Uniform manifold approximation projection (UMAP) of gene expression data from both COVID-19 patients. Cluster corresponds to the transcriptional clustering performed on the GEX datasets after excluding TCR and BCR receptor genes. Each point corresponds to a cell in one of the two patients. (B) Distribution of the fraction of cells located in each transcriptional cluster for the top 10 most expanded T cell clones of a single COVID-19 patient. Only those cells found in both GEX and VDJ sequencing datasets were included in the quantification. T cell clone was defined by unique CDRβ3 + CDRα3 nucleotide sequence. (C) The ten most expanded T cell clones defined by unique CDRβ3 + CDRα3 nucleotide sequence from a single COVID-19 patient are highlighted on the UMAP containing all cells from both patients. Each point corresponds to a unique cell barcode. Only those cells found in both GEX and VDJ sequencing datasets were included in the quantification.

Next we determined whether we could use Platypus to identify potentially virus-reactive B and T cell clones by integrating repertoire metrics such as somatic hypermutation and clonal similarity with phenotypic markers of activation and differentiation. We first noticed that 30 most expanded B cell clones had undergone somatic hypermutation in their VH segments, and since these patients were convalescent but still actively infected, highly mutated antibodies represent potential specific clones to SARS-CoV-2 (Figure 4A). Next, we used Platypus to infer the phylogenetic tree for the most expanded B cell clone while also annotating information about clonal expansion (based on identified number of cell barcodes) (Figure 4B). Surprisingly, we uncovered that 59 cells produced the exact same, full-length nucleotide antibody sequence and was actually the least mutated relative to the unmutated germline ancestral sequence (Figure 4B). The potential specificity of an antibody that has few somatic hypermutation is compatible with the recent discovery that B cells from COVID-19 patients that have germline-like antibodies with specificity to SARS-CoV-2 ^22^.

**Figure 4.**
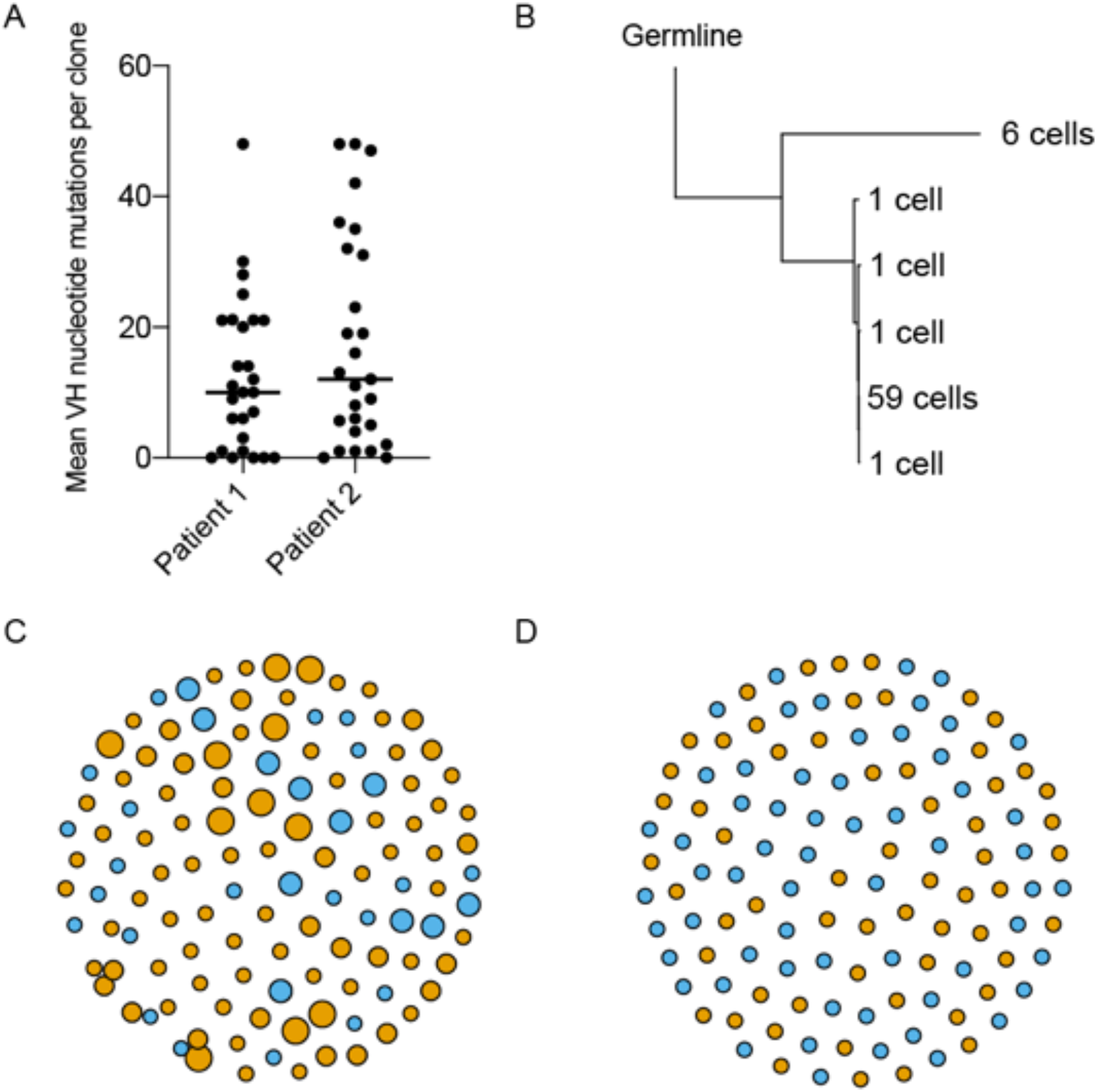
Internal functions from Platypus extract parameters relevant to the discovery of antigen-specific B and T cell clones. (A) Mean nucleotide somatic hypermutation for the 30 more expanded B cell clones found in the blood repertoire following SARS-CoV-2 infection. Somatic hypermutation was quantified in the V and J segments for both the heavy and light chains for each cell by comparing misalignments to the reference germline segments. (B) Phylogenetic tree rooted by germline reference sequence. The reference germline as determined by cellranger was set as the root. The number of cells in the tip label corresponds to the number of unique nucleotide variants producing the exact, full-length antibody sequence. Each tip label represents a single unique nucleotide paired V_H_ + V_L_ sequence. (C-D) Similarity networks depicting the B and T cell clones that are separated by less than 7 amino acid mutations in either CDRH3 or CDRL3 sequence (CDRβ3 + CDRα3 for T cells). Vertices represent unique clones defined as those cells containing identical CDHR3 + CDRL3 sequence (CDRβ3 + CDRα3 for T cells). Touching nodes connected by edges represent those clones that are separated by an edit distance of less than 7 amino acid mutations in their CDR3 regions.

We next questioned whether we could detect similar CDR3 sequences shared between the two COVID-19 patient repertoires by inferring sequence similarity networks for the top 60 most expanded B and T cell clones (Figures 4C, 4D) ^1^. This involved first calculating the edit distance between the CDR3s of clones from both patients and then drawing edges between those clones that were separated by less than 7 amino acid mutations in either the CDRH3 or the CDRL3 (or CDRβ3 + CDRα3 in T cells). This revealed that within B cell repertoires, but not T cell repertoires, there were B cell clones that were connected across patients, thereby suggesting convergence of antibody sequences which may be a result of specificity to SARS-CoV-2. Highly similar antibody sequences from the memory B cells of different COVID-19 patients have recently been shown to occur ^27^. The lack of connected T cell clones may be expected since it is likely these patients were not HLA-matched, in addition to the small portion of the blood repertoire sampled when performing single-cell sequencing.

We lastly investigated whether signatures of activation or differentiation would reveal potential B and T cell clones that may have recently interacted with viral antigen. Using B and T cell-specific gene sets contained internally in Platypus, we could investigate phenotypic and differentiation (ontogeny) markers for expanded clones (Figure 5). In the case of B cells, this included the following markers: B220 (PTPRC), MS4A1, CD27, CD38, and CD138 (SDC1), which distinguish between naïve, memory, and plasma cells subsets (Figure 5A). This analysis further supported a transcriptional heterogeneity within the most expanded clones, as demonstrated by individual cells expressing varying degrees of B cell defining markers such as CD38, XBP1, CXCR4, and TMSB10. Clones expressing high levels of such activation markers (CD38) or genes associated with plasma cell differentiation (XBP1) are potentially interesting candidates for virus-specific antibodies ^28^.

**Figure 5.**
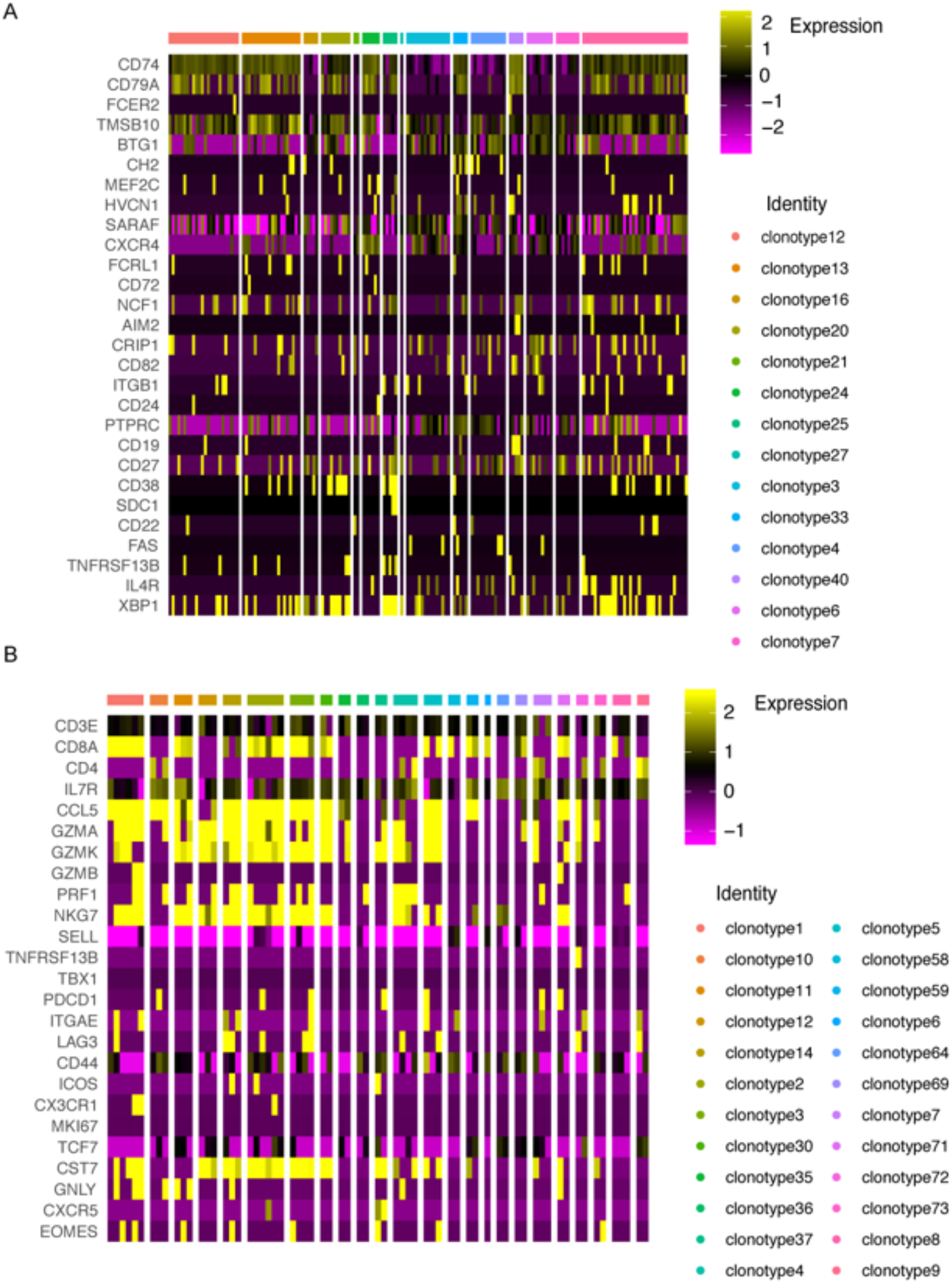
(A-B) Heatmap depicting normalized gene expression for the most expanded B and T cell clones, respectively, that were found in both the VDJ and gene expression (GEX) sequencing libraries from the blood repertoires of COVID-19 patients. The color of each column corresponds to an individual clonal family and the width of the bar corresponds to the number of cells found in the GEX library for that clone. Clone is defined as those cells containing identical CDHR3 + CDRL3 sequence (CDRβ3 + CDRα3 for T cells).

Despite the low levels of clonal expansion in the blood repertoire, we nevertheless questioned if cells belonging to the same clonal family demonstrated transcriptional heterogeneity based on commonly used T cell markers. Using the built-in gene sets internal to Platypus, we could visualize expression levels for genes such as CD44, PD-1 (PDCD1), Lag3, Tcf-7, granzymes, perforin, T-bet, Eomes, among others, at single-cell resolution (Figure 5B). Together, these genes allowed us to distinguish between naïve (CD44-, TCF1+, SELL+), memory (IL7R+), effector (KLRG1+), and exhausted (PDCD1+, LAG3+) subtypes that coexist within clonotypes (Figure 5B). Of interest was that the cells arising from expanded clones (e.g. clonotypes 1, 2, 3) expressed CCL5 and granzymes, whereas several clones corresponding to either one or two cells (clonotype 64, 69, 71, 72, 73) did not express these markers (Figure 5B). Platypus enables the user to relate clonotype information to user-defined transcriptional signatures, thereby connecting phenotypes involving antibody secretion, T cell exhaustion, amongst others to immune receptor sequence. Importantly, the user can customize the genes of interest to explore in the context of expanded clones. Taken together, Platypus enabled us to investigate and visualize repertoire and transcriptome parameters that can help facilitate the rapid discovery of relevant B and T cell clones.

## Discussion

As in many areas of biology, researchers in adaptive immunity are poised to heavily utilize scSeq as an essential method to address basic and translational questions, therefore creating a substantial need for suitable bioinformatics tools. We have demonstrated here that Platypus enables rapid extraction, integration, and analysis of scSeq of lymphocyte repertoires and transcriptomes to uncover meaningful insight on lymphocyte immunobiology and function. By using convalescent COVID-19 patients as an example, we were able to detect clonally expanded B and T cells at the single-cell resolution from less than 10,000 cells, suggesting these clones are present at high-levels in the remainder of the blood repertoire. Interestingly, we observed many expanded clones were of the IgA isotype and often did not coexist with other isotypes, highlighting that IgA antibodies may play an important role in the response to SARS-CoV-2. We could relate these expanded clones to diverse cellular phenotypes based on district transcriptional cluster groups, which revealed that even when cells belonged to the same BCR or TCR clonal group, surprisingly they could still be found in different transcriptional-clusters. This highlights that lymphocytes sharing the same immune receptor specificity may still undergo very different cell fates and function in the course of an immune response. We could furthermore analyze various repertoire parameters including somatic hypermutation, clonal evolution, and clonal convergence at the single-cell resolution. Platypus allowed us to analyze and extract full-length, paired, heavy-light chain information for clonally expanded and heavily mutated clones, which offers a possible approach to identify SARS-CoV-2-specific lymphocytes. Leveraging Platypus, we were able to visualize this information and have full-length BCR and TCR sequences ready for cloning in less than 10 lines of code – something that can greatly accelerate the discovery of adaptive immune therapeutics. In conclusion, Platypus enables a broad range of immunologists and bioinformaticians alike to gain quantitative insight at a single-cell resolution of how immune repertoire parameters relate to heterogeneous transcriptome information.

## Materials and Methods

### Patient samples

Patients were participants of the SERO-BL-COVID-19 study sponsored by the Department of Health, Canton Basel-Landschaft, Switzerland. Both patients tested positive for SARS-CoV-2 after RT-PCR of naso- and oropharyngeal swab and did not require hospitalization. Whole blood was collected 31 and 32 days following a positive RT-PCR test and subjected to density gradient centrifugation using the Ficoll Paque Plus reagent (GE Healthcare, #17-1440-02). After separation, the upper plasma layer was collected for ELISA detection of IgG and IgA SARS-CoV-2-specific antibodies (Euroimmun Medizinische Labordiagnostika, #EI2668-9601G, #EI2606-9601A). Peripheral blood mononuclear cells (PBMC) were collected from the interphase, resuspended in freezing medium (RPMI 1640, 10%(v/v) FBS, 10%(v/v) dimethyl sulfoxide) and cryopreserved in liquid nitrogen. Point-of-care lateral flow immunoassays assessing the presence of IgG and IgM SARS-CoV-2-specific antibodies (Qingdao Hightop Biotech, #H100) were performed at the time of blood collection.

### Immunomagnetic isolation of B cells and T cells

PBMC samples were thawed, washed in complete media (RPMI 1640, 10%(v/v) FBS) and pelleted by centrifugation. Cells were resuspended in 0.5 mL complete media, counted and treated with 10 U ml^−1^ DNAse I (Stemcell Technologies, #) for 15 min at RT in order to prevent cell clumping. After DNase I digestion, cells were washed in complete media, pelleted by centrifugation and resuspended in 0.5 mL flow cytometry buffer (PBS, 2%(v/v) FBS, 2 mM EDTA). The cell suspension was filtered through a 40 μM cell strainer prior to immunomagnetic isolation. As a first step, plasma cells were isolated using the EasySep Human CD138 Positive Selection Kit II for future studies (Stemcell Technologies, #17877). The negative fraction of the above selections was divided into two aliquots that were subjected to negative immunomagnetic isolation of either B cells (EasySep Human Pan-B cell Enrichment Kit, Stemcell Technologies, #19554) or T cells (EasySep Human T cell Isolation Kit, Stemcell Technologies, #17951). After isolation, B cells and T cells were pelleted by centrifugation, resuspended in PBS, 0.4% BSA(v/v), filtered through a 40 μM cell strainer and counted. T cells and B cells originating from the same patient were pooled in equal numbers and the final suspension was counted and assessed for viability using a fluorescent cell counter (Cellometer Spectrum, Nexcelom). Whenever possible, cells were adjusted to a concentration of 1×10^6^ live cells/mL in PBS, 0.04%(v/v) BSA before proceeding with droplet generation.

### Single cell sequencing libraries

Single cell 10X libraries were constructed from the isolated single cells following the Chromium Single Cell V(D)J Reagent Kits User Guide (CG000086 Rev M). Briefly, single cells were co-encapsulated with gel beads (10X Genomics, 1000006) in droplets using 4 lanes of one Chromium Single Cell A Chip (10X Genomics, 1000009). V(D)J library construction was carried out using the Chromium Single Cell 5’ Library Kit (10X Genomics, 1000006) and the Chromium Single Cell V(D)J Enrichment Kit, Human B Cell (10X Genomics) and Human T Cell (10X Genomics). The reverse transcribed cDNA was split in three and GEX, B and T cell V(D)J libraries were constructed following the instructions from the manufacturer. Final V(D)J libraries were pooled and sequenced on the Illumina NovaSeq platform (300 cycles, paired-end reads). Pooled 5’ gene expression libraries were and sequenced on the Illumina NextSeq 500 (26/91 cycles, paired-end) with a concentration of 1.6 pM with 1% PhiX. Resulting FASTQ files were demultiplexed and subsequently used as input to cellranger (v3.1.0, 10x Genomics). GEX sequencing libraries were aligned to the refdata-cellranger-GRCh38-3.0.0 reference genome and VDJ genes and the VDJ sequencing libraries were aligned to the vdj_GRCh38_alts_ensembl-3.1.0-3.1.0 reference using Single Cell V(D)J R2-only chemistry.

### Immune repertoire analysis using Platypus

The R package, accompanying code, and processed sequencing data used in this study are publicly available at github.com/alexyermanos/Platypus and doi.org/10.5281/zenodo.4140161. Briefly, clonotyping information was extracted directly from the output directory of cellranger using the function analyze_VDJ in Platypus. Quantifying the number of unique clones was performed using the VDJ_clonotype function in Platypus, with clone.strategy set to either “cdr3.aa”, “hvj.lvj”, hvj.lvj.cdr3length.cdr3homology”, or “hvj.lvj.cdr3lengths”. The isotype distributions for the top thirty B cell clones were calculated using the VDJ_isotypes_per_clone function in Platypus. CDR3 length distribution and sequence logo plots were calculated on the output of the VDJ_per_clone function. Sequence logos were calculated based on the function ggseqlogo from the R package ggseqlogo, with method set to “prob” and seq_type set to “aa”. The output directory from cellranger count was supplied as input to the function automate_GEX, which analyzes and integrates transcription data using functions from the R package Seurat. Briefly, the GEX libraries were integrated using the SCTransform function from Seurat. Cells containing more than 20% mitochondrial genes were removed. TCR and BCR genes were filtered prior to integration and gene expression analysis. The number of variable features selected was 2000 for the RunPCA function using the first 10 dimensions and Cluster resolution was set to the default 0.5. Transcriptional cluster and clonotype information were integrated using the VDJ_GEX_integrate function in Platypus under default parameters. Quantification of the transcriptional cluster distribution for the 10 most expanded clones from patient 1 were calculated and visualised using the VDJ_GEX_expansion function in Platypus. Those cells from the two most expanded clones containing barcodes in both VDJ and GEX datasets were highlighted on the UMAP plot using the function visualize_clones_GEX in Platypus under default parameters. Somatic hypermutation was calculated in the VDJRegion as defined by MiXCR via the function call_MIXCR in Platypus, which utilized MiXCR (v3.0.12) ^22^. The number of nucleotide alignment mismatches between the reference germline and the full-length VDJRegion for both heavy and light chain nucleotide sequence was then computed based on the best alignment determined by MIXCR. The phylogenetic tree was inferred by appending the full-length VDJRegion of the heavy and light chain for each unique sequence after appending VH VDJRegion and VL VDJRegion, as determined from the output function of call_MIXCR in Platypus. The reference germline sequence was first extracted from the initial cellranger alignment using the function VDJ_extract_germline in Platypus and added to the set of input sequences which were supplied to VDJ_tree. Sequence similarity networks were calculated using the function VDJ_networks in Platypus by calculating the edit distance separately for CDRH3 and CDRL3 amino acid sequences and then summing the two matrices. Edges were then drawn between those clones separated by less than 7 amino acid mutations. The heatmap integrating clonotype membership with user-defined gene lists was performed using the GEX_heatmap function in Platypus.

## Acknowledgements

We acknowledge and thank Dr. Christian Beisel, Elodie Burcklen and Ina Nissan, at the ETH Zurich D-BSSE Genomics Facility Basel for excellent support and assistance.

## Funding

This work was supported by the European Research Council Starting Grant 679403 (to STR) and ETH Zurich Research Grants (to STR and AO).

## Author Contributions

AY, AG, JY, CP, DR, FB, RVL, and DN performed experiments and analyses. All authors contributed to the study and manuscript design.

## Competing Interests

There are no competing interests.

## Data and Materials Availability

The R package, code, and example data used in this publication are found at github.com/alexyermanos/Platypus and https://doi.org/10.5281/zenodo.4140161.

---

**Figure S1.**
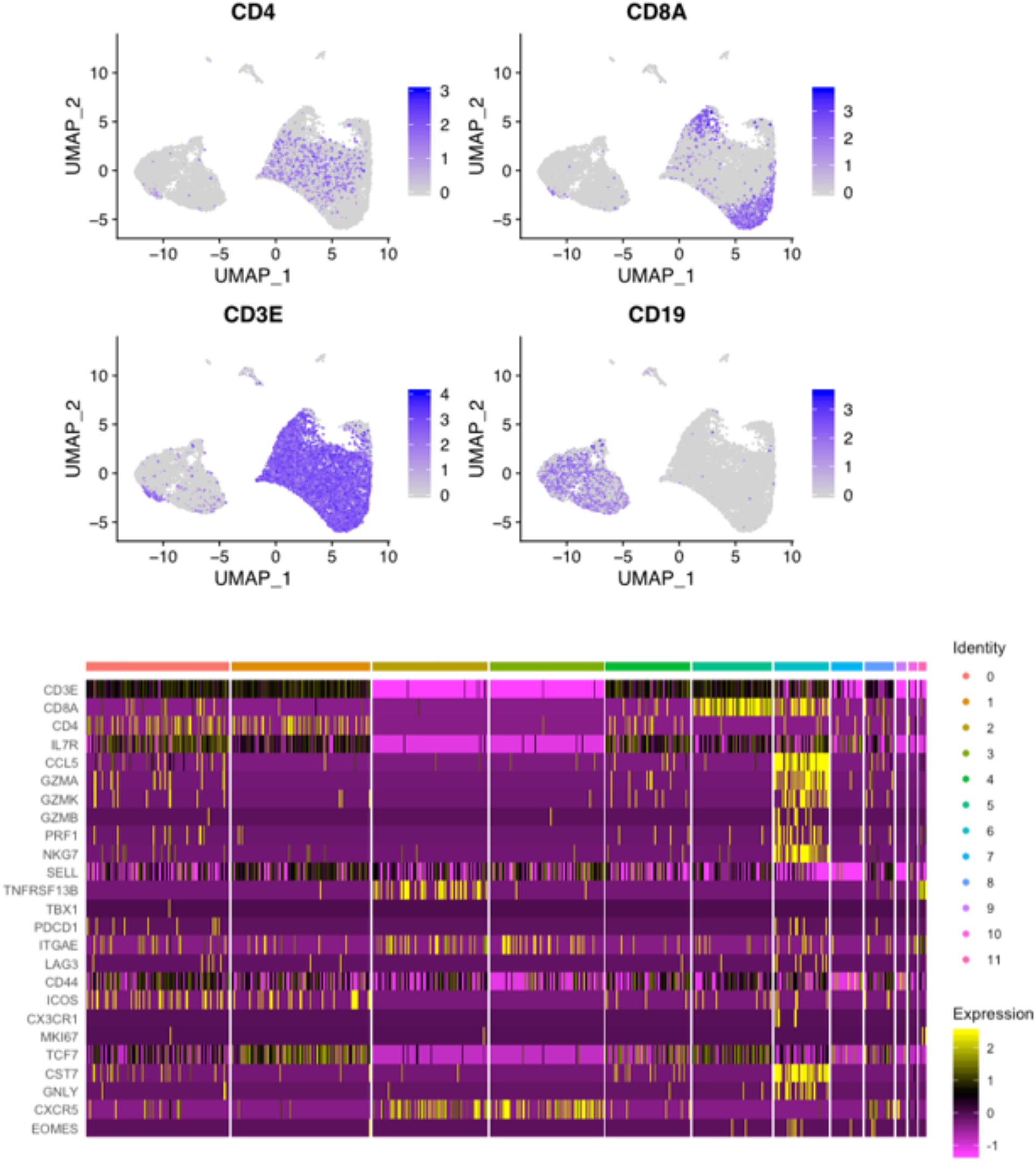
Expression of selected B and T cell genes. (A) Uniform manifold approximation projection (UMAP) plot depicting the expression of CD4, CD8A, CD3E, and CD19 to identify B and T cells. (B) Normalized expression of common T cell genes distributed by cluster. Each column represents a single cell and column identity refers to transcriptional cluster as previously determined (Figure 3A). The width of the column identity corresponds to the number of cells within each cluster.

## Notes

https://doi.org/10.5281/zenodo.4140161

https://github.com/alexyermanos/Platypus

### Competing Interest Statement

The authors have declared no competing interest.

